# Extracellular Lactate Inhibits HIV Infection in Dendritic Cells by Downregulating CD4 Expression

**DOI:** 10.64898/2026.11.13.766458

**Authors:** Mayra Marquez, Qian Chen, Whitney Bruchey, Jun Yang, Udeshika Kariyawasam, Jordan Metz, Bharatwaj Sowrirajan, Adam Rupert, Sylvain Laverdure, Tomozumi Imamichi

## Abstract

Cellular metabolism and cellular microenvironments influences immune-cell function. We previously demonstrated that interleukin-27 (IL-27)-induced monocyte-derived dendritic cells (27DCs) exhibit resistance to HIV-1. A mechanistic study for this antiviral effect suggested that in addition to inducing antiviral genes, IL-27 may influence cellular metabolism that contributes to the anti-HIV effect. During course of study, we observed that 27DCs accumulated lactate in culture supernatants by 5.7-fold increase compared with cultures of untreated DCs (iDCs) (12.1±1.3 versus 2.1±0.2 mM). Here, we assesses extracellular role of lactate on HIV infection. Healthy donors’ CD14+ monocytes were differentiated to DCs using GM-CSF and IL-4 in the absence of presence of lactate (Lac). DCs were infected with HIVAD8. Lactate-induced DC (LacDC) reduced R5-tropic HIV-1 AD8 replication by 71.1±8.3% without detectable cytotoxicity compared to that in untreated cells (iDCs). LacDCs exhibited reductions in viral particles binding, membrane fusion of viral particles, and HIV reverse-transcription by 78.5±2.8% (n=3, p<0.05), 78.7±3.3% (n=3, p<0.01), 86.2±5.7% (n=3. p<0.01), respectively. FACS analyses demonstrated that surface CD4 expression on LacDC decreased by 56.6±5.1% (n=5, p<0.05) compared to iDC, whereas total cellular CD4 protein amount and surface CCR5 expression were not significantly altered. Neither the lactate transporter inhibitor nor the lactate receptor restored HIV-1 susceptibility. These findings demonstrate that lactate exposure during dendritic-cell differentiation induces an HIV-1-resistant phenotype associated with reduced surface CD4 availability and impaired viral attachment. This study identifies extracellular lactate as a modulator of dendritic-cell susceptibility to HIV-1 and suggests a potential link between IL-27-associated metabolic changes and antiviral immunity.

## Introduction

In recent years, a substantial number of studies have been conducted in the area of immunometabolism, examining how changes in intracellular metabolic pathways influence immune cell function [1,2]. Immunometabolism is closely linked to differentiation of dendritic cells [2-4]. In glycolysis, glucose is split into two molecules of pyruvate, and under lactic acid fermentation, pyruvate is converted into lactate [5,6]. Accumulated lactate in cytoplasm effluxes from cells through Monocarboxylate Transporters (MCTs: MCT1 and MCT4). In addition, extracellular lactate can influence immune cells through signaling mediated by the lactate receptor GPR81 (also known as hydroxycarboxylic acid receptor 1: HCAR1), or GPR132 (also known as GA2) [7]. In plasmacytoid DCs, lactate suppresses type I interferon production through mechanisms involving both GPR81 and intracellular lactate uptake [8], which may influence viral infection in DC or microenvironments.

We have investigated the mechanism underlying resistance of HIV infection in interleukin-27 (IL-27)-induced monocyte-derived dendritic cells (27DC) [9]. Although we obtained gene expression profiles, difficulties with siRNA transfection in DCs prevented identifying a key factor regulating HIV inhibition through siRNA-mediated knockdown assays [9,10]. During further characterization of 27DC, we observed increased lactate accumulation in culture supernatants of 27DC. This observation prompted us to investigate whether lactate itself could influence HIV infection. In the current study, we report the potential role of Lactate on HIV infection.

## Materials and Methods

### Cells and reagents

Immature dendritic cells (iDCs) were generated from CD14+ monocytes from healthy as previously described [9,10]. IL-27- or lactate-induced DC (LacDC) were generated by supplement of 100 ng/ml of IL-27 (R&D Systems, Minneapolis, MN, USA) or 31.6 mM sodium lactate (Sigma-Aldrich, St. Louis, Missouri) in the culture of monocytes. Macrophage (M-Mac) or IL-27-differentatied macrophages (I-Mac) were prepared as previously described (Ref). AZD3965 and Syrosingopine were obtained from MedChemExpress (Monmouth Junction, NJ). Cell viability was measured using the Nexcelom Cellometer Auto 2000 (Revvity, Waltham MA, USA) with by AO/PI exclusion staining dye (Revvity).

### Measurement of Extracellular pH and lactate concentration

Cell-free culture supernatants from iDCs and 27DCs were collected after differentiation. The pH in the culture media was measured using a pH meter (Corning, Corning, NY), and lactate concentrations were measured using the Lactate-Glo Assay (Promega, Madison, WI, USA) according to the manufacturer’s instructions.

### Qualification of ATP concentration

Intracellular ATP in iDC and 27DC was quantified using CellTiter-Glo Luminescent Cell Viability Assay kit (Promega) according to the vendor protocol, ATP amounts were measured using a Tecan Spark multimode microplate reader. Concentration of ATP was determined by using a standard ATP (Sigma Aldrich). Total cellular protein was determined using a bicinchoninic acid (BCA) protein assay kit (Thermo Fisher Scientific).

### Virus Production and Quantification

Infectious HIVAD8 was generated as described previously [11]. For HIV-1 fusion assays, HIVNLAD8-ΔVpr virions containing Vpr-BlaM (HIV-BlaM) were generated as previously described [12,13]. Infectious viral titers were determined as the 50% tissue culture infectious dose (TCID50) using an endpoint dilution assay with the HIV-1 p24 ELISA Kit (Revvity), as previously described [14].

### HIV-1 replication assay

iDCs and LacDCs (1×10^6^ cells) were infected with HIVAD8 (5,000 TCID50) as previously described [9]. After infection, cells were cultured for 14 days. HIV-1 released into the supernatants was measured using the HIV-1 p24 ELISA Kit [9].

### HIV-1 Binding Assay

HIVAD8 binding assay was conducted as described previously [9]. iDCs or LacDCs were incubated with HIVAD8 for 2 h at 4°C. Following incubation, cells were washed and lysed in 100 µL radioimmunoprecipitation assay (RIPA) buffer (Boston BioProducts, Milford, MA, USA) supplemented with protease, EDTA and phosphatase inhibitor cocktails (Thermo Fisher Scientific). Cell-associated HIV-1 was quantified by measuring p24 antigen using the HIV-1 p24 ELISA Kit. Total cellular protein concentration was determined using a BCA assay kit, and p24 levels were normalized to total protein content and expressed as picograms of p24 per microgram of total protein (pg p24/µg total protein) [9].

### Quantitative Reverse Transcription PCR (qRT-PCR)

Quantitative reverse transcription PCR (qRT-PCR) was performed using 1 µg of total RNA with TaqMan Gene Expression Assays (Thermo Fisher Scientific) specific for CD4, CCR5, or GAPDH. as previously described [9]. Reactions were performed using an iQ5 Real-Time PCR Detection System (Bio-Rad, Hercules, CA, USA), as previously described [9].

### HIV-1 Fusion Assay

Fusion of HIV-1 particles with iDCs and LacDCs membrane was assessed using HIV-BlaM as previously described [9,12] with minor modification. Cells were incubated with 500 ng/ml of HIV-BlaM and then loaded with 1× CCF2-AM fluorescent substrate using the β-lactamase loading kit (Invitrogen). Cleaved CCF2-AM substrate by fusion were analyzed using an Amnis imaging flow cytometer (Cytek Biosciences, Fremont, California) HIV-1 fusion was quantified based on CCF2-AM substrate cleavage [9,12].

### HIV-1 Late Reverse Transcription Products

HIV-1 late reverse transcription (RT) products were quantified using 100 ng of genomic DNA from HIV infected cells as previously described [9]. HIV-1 late RT product copy numbers were normalized to RNaseP [9].

### Western Blotting

Western Blotting (WB) were conducted as previously described [9]. Antibodies used in WB is listed in supplemental table 1. Band intensities were quantified by densitometry using ImageJ (NIH), as previously described[15]. β-actin was used as the loading control, and the expression levels of MCT1, MCT4, CD4, and CCR5 were normalized to β-actin.

### Flow Cytometric Analysis of CD4 and CCR5 Surface Expression

Surface expression of CD4 and CCR5 on cells were assessed by flow cytometry as previously described [9]. Antibodies used in the assay is listed in Supplementary table 1. The results were analyzed using FCS Express version 7 (DeNovo Software, Pasadena, CA, USA).

### Construction of RNA-Seq libraries and gene expression profile data analysis

Gene expression profile was analyzed using bulk RNA seq data. The RNA seq library was constructed using stranded RNA seq kit (Illumina, San Diego, CA, USA), and sequence was run on NextSeq 2000 (Illumina), data was analyzed using Partek Flow software. Detail method is described in supplemental methods.

### Statistical Analysis

Intergroup comparisons were performed using one-way ANOVA or *t-*test (GraphPad, San Diego, CA, USA). p-values <0.05 were considered statistically significant: * p < 0.05, ** p < 0.01, *** p < 0.001, **** p < 0.0001; p ≥ 0.05 was considered not significant (ns).

## Results

### 27DC exhibited extracellular acidification and increased lactate accumulation

We previously demonstrated that IL-27-induced dendritic cells (27DCs) exhibit potent resistance to HIV-1 infection. During differentiation, we observed a distinct change in the color of the culture medium from 27DCs compared with immature dendritic cells (iDCs) **(Figure 1A)**. This significant change was not observed in M-CSF-induced macrophage (M-Mac) and IL-27-indcued macrophages (I-Mac) [16]. Measurement of pH in culture-medium of iDCs and 27DCs exhibited pH=7.46±0.04 (n=3) and pH=7.04±0.05 (n=3), respectively; demonstrating a significant acidification in 27DCs (p<0.01; **Figure 1B)**. We hypothesized that the increased proton concentration ([H⁺]) in the culture medium might involve ATP-dependent proton transport across the plasma membrane [17]. We therefore measured intracellular ATP levels, which were comparable between iDCs and 27DCs (**Figure 1C**). Thus, the observed difference in extracellular pH was not accompanied by a detectable difference in intracellular ATP levels. Another potential mechanism for the increase in [H⁺] was the export of lactate through proton-coupled monocarboxylate transporters (MCT1 and MCT4), since this process is accompanied by H⁺ release. Quantification of the extracellular lactate assay demonstrated that the lactate concentration in 27DC increased by 5.8-fold compared with that in iDC (p<0.001 see **Figure 1D)**. This indicated that the increase in proton concentration was most likely due to lactate accumulation. Notably, lactate concentration was not changed in IL-27-induced Macrophages (I-Mac), indicating that lactate accumulation is unique function in DC culture.

**Figure 1.**
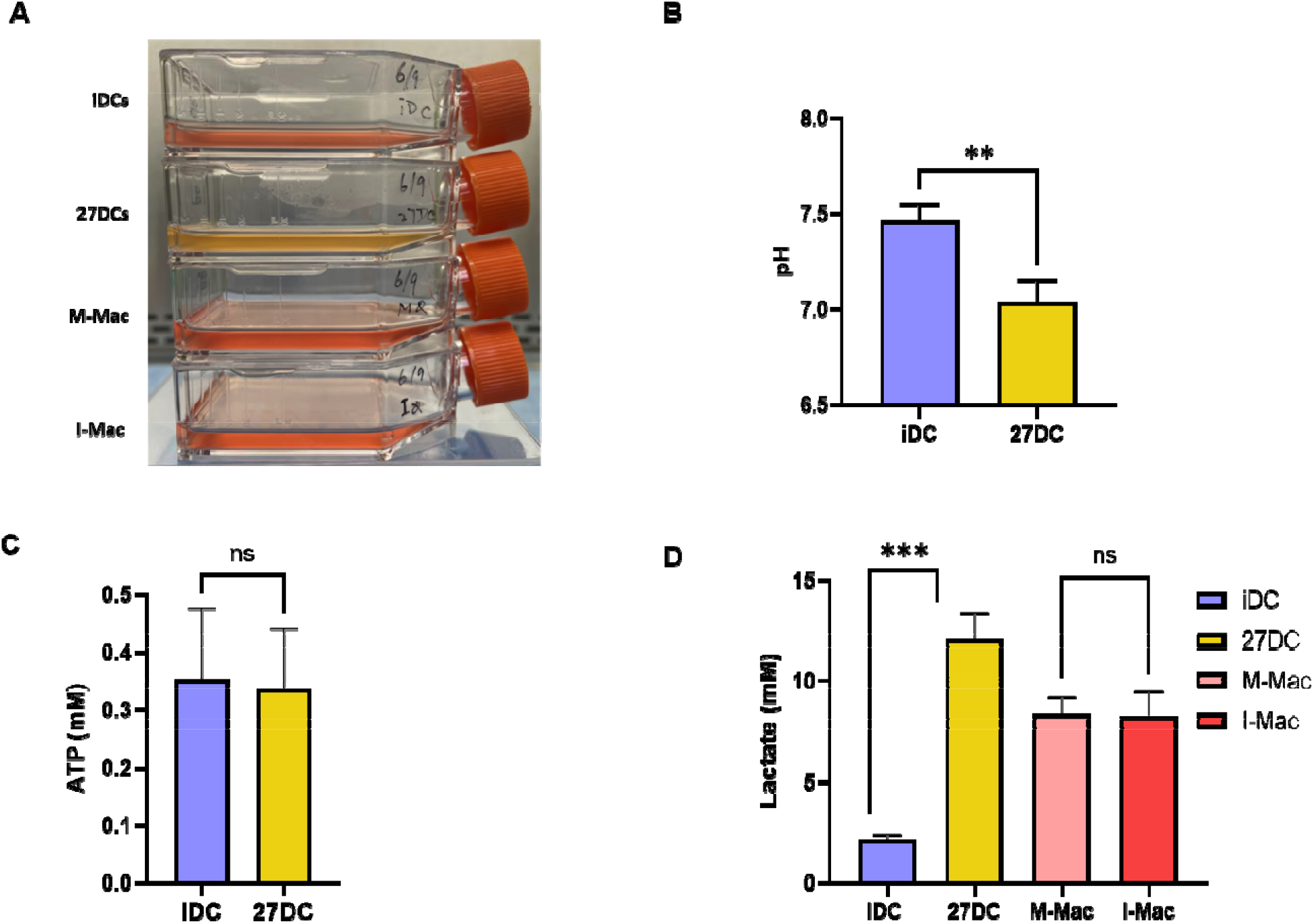
Acidification and extracellular lactate accumulation induce in 27DC culture. **(A)** Representative image of iDC, 27DC, M-Mac and I-Mac cultures. **(B)** Culture-medium pH was measured in iDC and 27DC cultures on day 7 of differentiation by pH meter **(C)** ATP concentrations in cells were measured using CellTiter-Glo Luminescent Cell Viability Assay kit as described in Materials and Methods. **(D)** Extracellular lactate concentrations were measured in cell-free culture supernatants from iDCs, 27DCs, M-Mac and I-Mac on day 7 using the Lactate-Glo assay kit. Data represents the mean ± SE (n=7 for iDC and 27DC, n=4 for M-Mac and I-Mac). t-test was conducted ** p < 0.01, *** p < 0.001, ns: not significant.

### Lactate differentiates monocytes to HIV-1-resistant dendritic-cells

To determine whether extracellular lactate contributes to differentiation into HIV-resistant DC, CD14+ monocytes were differentiated into iDCs in culture medium with different concentrations of lactate. As shown **Figure 2A**, lactate induced HIV resistant DC dose dependent manner and reached maximal activity at 31.6mM without cytotoxic effect (iDC 86.7±1.5% and LacDC 87.2±2.1%). Of interest, lactate had no impact on macrophage differentiation, thus lactate effect is cell type dependent. The dendritic cells differentiated in the presence of 31.6 mM sodium lactate (LacDCs) suppressed HIV replication by 71.1±8.3% (n=11) compared with iDCs (p<0.05, **Figure 2B**). Thus, exposure to lactate during DC differentiation was sufficient to generate resistance to HIV1. We next investigated individual steps of HIV-1 replication cycle in LacDCs.

**Figure 2.**
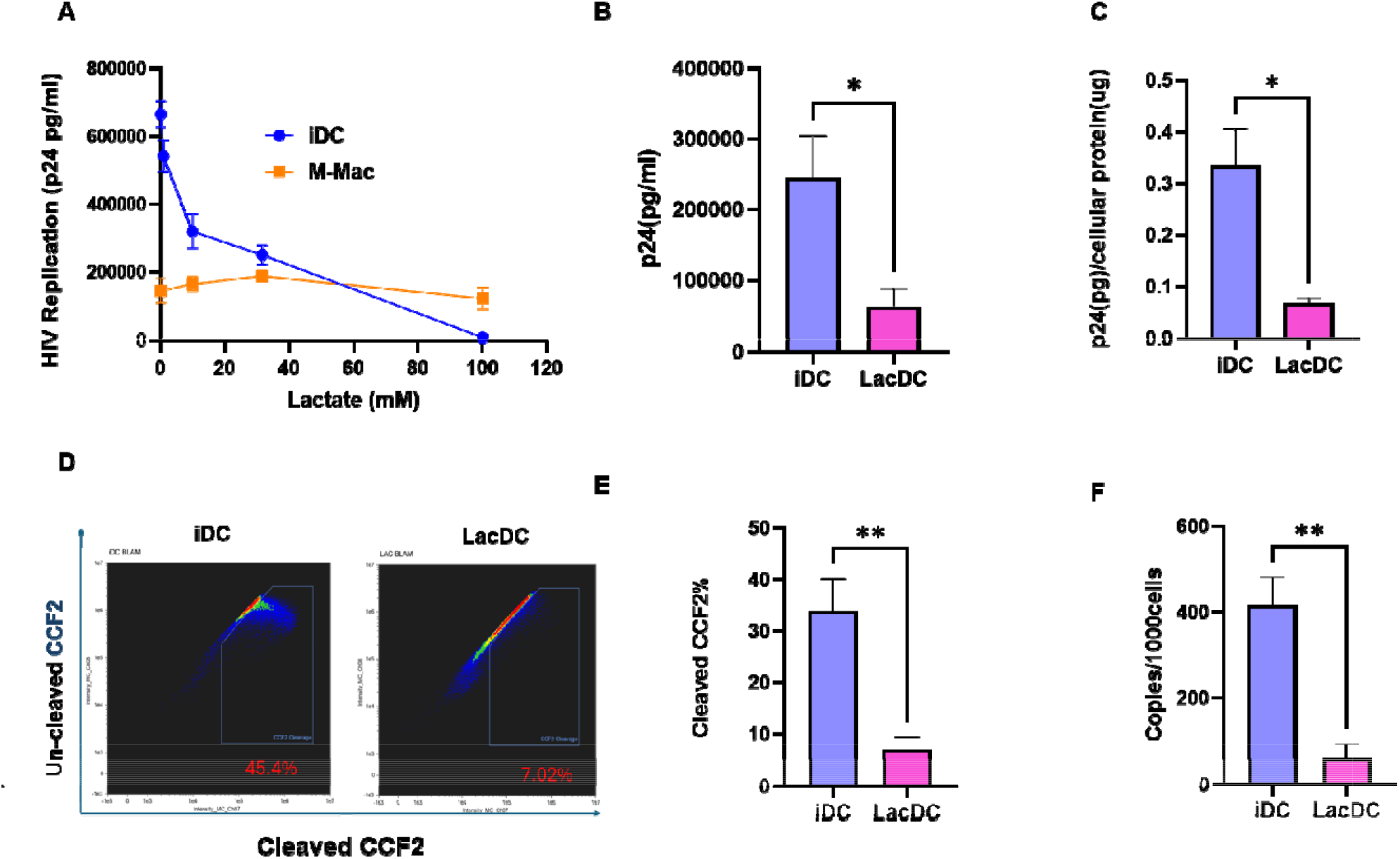
Lactate Restricts HIV-1 Binding, viral fusion, and reverse transcription. **(A)** iDC and M-Mac were induced in the presences of different dosage of lactate, and then infected with HIV. (B) 27DCs was induced in the presence and absence of 31.6 mM Lac (n=11). Cells were infected with HIVAD8 and HIV replication was assessed using p24 Elisa kit. Data represents the mean ± SE. **(C)** HIV-1 binding assay was performed as described in Materials and Methods; the results of HIV-1 binding was expressed as pg p24/µg total protein. **(D and E)** iDCs and LacDCs were incubated with HIVAD8 virions containing BlaM-Vpr and subsequently loaded with the fluorescent substrate CCF2-AM. Viral fusion was detected by BlaM-mediated cleavage of CCF2 as described in Materials and Methods. Representative AMNIS dot plots are shown for one of six donors **(D)**. Summarized fusion assay result represents the mean ± SE (n=6) **(E)**. **(F)** HIV-1 late RT products in HIVAD8-infeced iDCs and LacDCs were quantified by real-time PCR as described in Materials and Methods. Result shows mean ± SE (n=3). The t-test was conducted. * p < 0.05, ** p < 0.01.

We first examined viral attachment. HIV-1AD8 binding was assessed at 4°C to permit viral attachment while preventing membrane fusion. Compared with iDCs, cell-associated HIV-1 was reduced by approximately 78.5±2.8% in LacDCs (n=3, p<0.05; **Figure 2C)**. We next assessed HIV-1 membrane fusion using a β-lactamase-Vpr (BlaM-Vpr) fusion assay. HIV-1 fusion was reduced by approximately 78.7±3.3% in LacDCs compared with iDCs (n=3, p<0.01; **Figure 2D and 2E**). To further define whether HIV-1 inhibition was restricted at reverse transcription (RT) step, after fusion, late reverse products were quantified 24 hours after HIV-1 infection. Late RT product copy numbers were reduced by approximately 86.2±5.7% in LacDCs compared with iDCs (n=3, p<0.01; **Figure. 2F**). Collectively, these findings demonstrated that LacDCs exhibited restriction at early binding stages with sustained suppression through fusion and late RT products; however, no additional suppression was observed.

### Lactate reduces cell-surface CD4 expression without altering CCR5

Given the marked reduction in HIVAD8 binding observed in LacDCs, we examined whether lactate altered the cell-surface expression of the HIV receptor (CD4 and coreceptor CCR5). Flow cytometric analysis illustrated that mean fluorescence intensity (MFI) of CD4 was reduced by approximately 56.6±5.1% in LacDCs compared with iDCs (n=5, p<0.05; **Figure 3A and 3B)**. In contrast, CCR5 surface expression was not significantly altered in either LacDCs compared with iDCs (**Figure 3A and 3B).**

**Figure 3.**
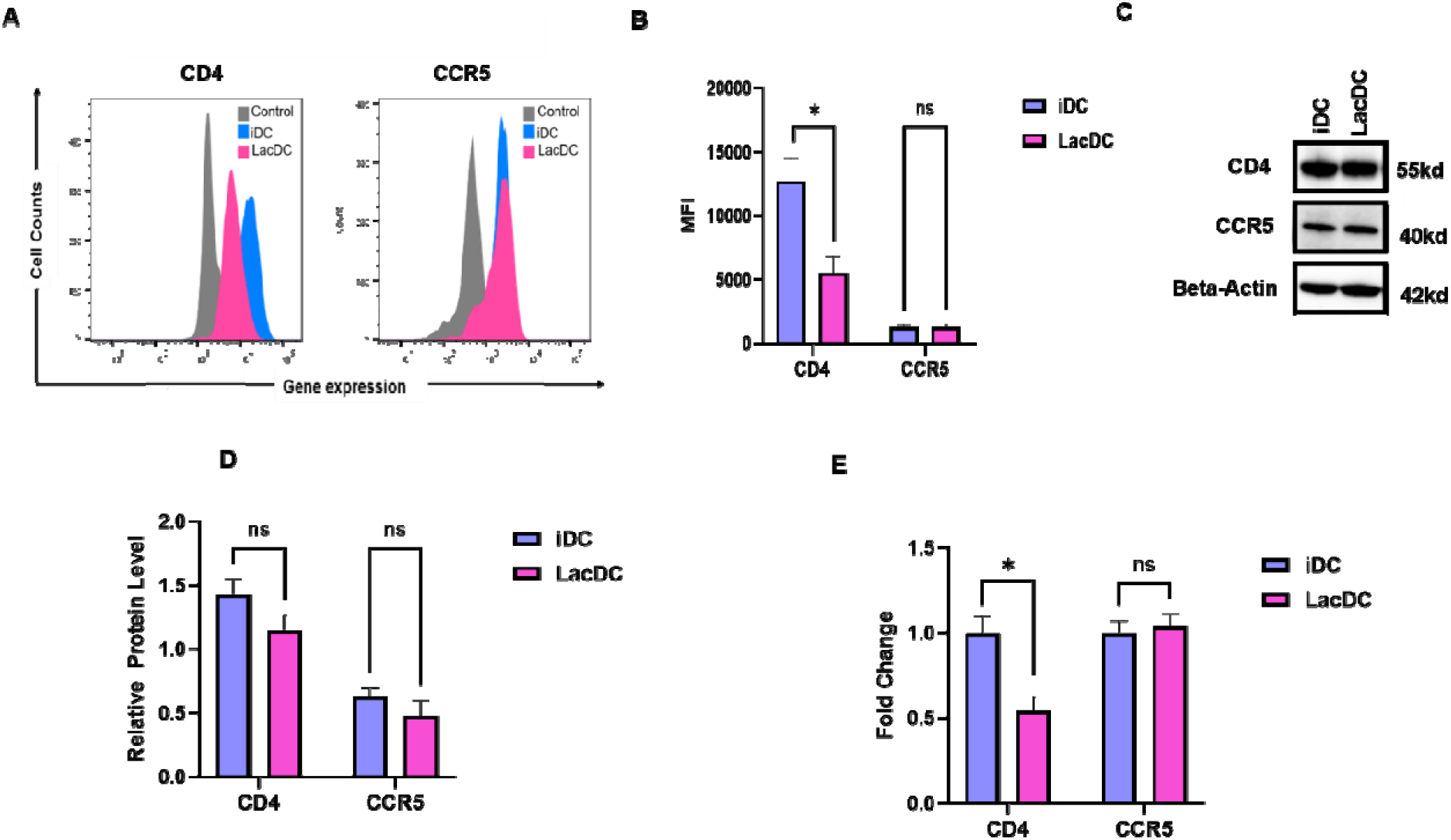
Lactate reduces CD4 expression. **(A)** Cell-surface CD4 and CCR5 was analyzed by flow cytometry as described in Materials and Methods. Gray shows isotype control, and pink and blue indicate iDC and LacDC, respectively. Representative data show from one donor of five donors. **(B)** Summary data are presented as means ± SE (n=5). **(C)** WB was performed using anti-CD4, anti-CCR5, and anti-beta-actin antibodies. A representative result shown. (**D**) The band intensity of each protein in WB was normalized by the band intensity of beta-actin. summarized WB results show, mean ± SE (n=3). **(E)** The qRT-PCR was conducted to define the expression level of each gene of interest using total cellular RNA from three independent donors. Data show mean ± SE (n=3). The T-test was conducted, * p < 0.05, ns; not significant.

To determine whether the reduction in surface CD4 was accompanied by a degradation of total cellular CD4 protein, CD4 and CCR5 expression was assessed by Western blot. Despite reduction in CD4 cell-surface expression, amount of cellular CD4 protein was not significantly altered in LacDCs compared with iDCs, while CCR5 expression was not changed (**Figure 3C and 3D)**. We subsequently examined CD4 and CCR5 transcript levels in iDC and LacDC using qRT-PCR. CD4 transcript levels were reduced by approximately 45.5±8.3% (n=3, p<0.05) in LacDCs compared with iDCs, whereas CCR5 transcript levels were not significantly altered (**Figure 3E)**.

### Extracellular lactate, not intracellular, induces anti-HIV effect

Given that LacDCs resisted HIV infection, we investigated whether lactate transporters (MCT1 and MCT4) contributes to the differentiation. Primary role of MCT1 and MCT4 is lactate uptake and lactate export, respectively [18]. Expression of MCT1 and MCT4 in iDCs and LacDCs was first assessed by WB. Although MCT1 and MCT4 expression demonstrated donor dependency among three donors (**Figure 4A)**. Overall, quantitative analysis revealed no significant differences in MCT1 or MCT4 expression between iDCs and LacDCs (**Figure 4B)**.

**Figure 4.**
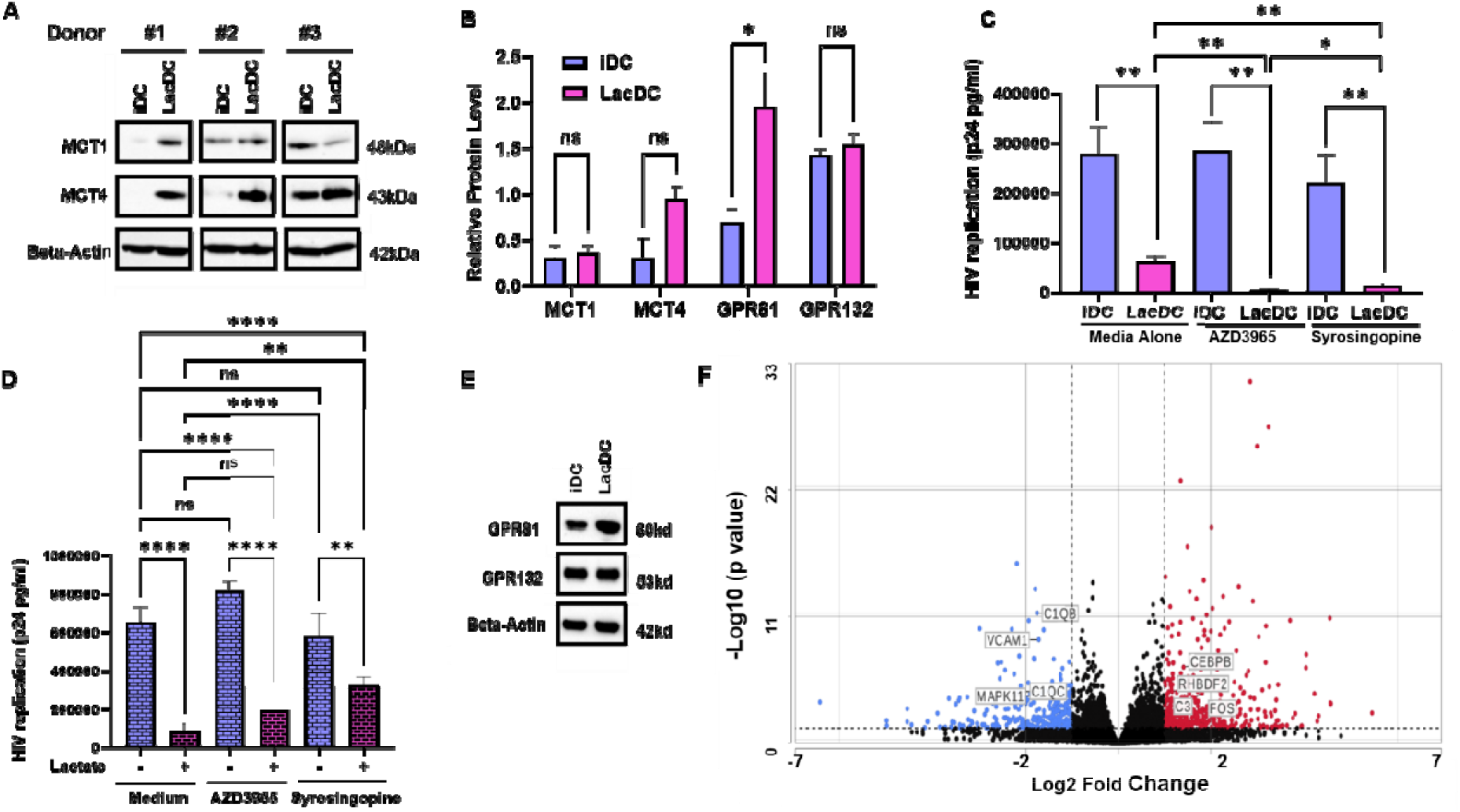
Role of Lactate transporters. **(A)** WB was performed using iDC and LacDC cell lysate from three independent donors. **(B)** The band intensity of each protein in WB was normalized by the band intensity of beta-actin and summarized. Results show, mean ± SE (n=3). **(C)** iDC or LacDC were induced in the absence or presence of 10 μM AZD3965 or 10 μM Syrosingopine. Cells were infected with HIVAD8 and then cultured for 14-days without inhibitors. HIV replication was quantified by p24 antigen ELISA. **(D)** iDC were infected with HIVAD8 and then cultured for 14 days in the absence or presence of lactate with or without AZD3965 or Syrosingopine. HIV replication was quantified by p24 antigen ELISA. **(E)** GPR81 and GRP132 expression in iDC and LacDCs was analyzed by WB as described in Materials and Methods. One of representative result from three independent donors is shown. **(F)** The gene expression profile was compared between lactate treated for one-day and 4 days from three independent donors using bulk RNA seq. To visualize the overall distribution of DEGs; a two-dimensional volcano plot was generated. The statistical significance threshold was set as p-value <0.05 and absolute fold change>=2. Genes exceeding these combined criteria were color-coded, significantly up-regulated in red and significantly down-regulated genes in blue, while non-significant genes remained black dots. Genes were extracted from KEEG pathway analysis through DAVID were labeled with its name.

To assess the contribution of lactate influx on anti-HIV effect, inhibitors targeting MCT1 or MCT4 were supplemented in the culture during inducing iDC or LacDC. Neither 10 µM AZD3965 (an MCT1 inhibitor) nor 10 µM syrosingopine (an MCT1 and MCT4 inhibitor) restored HIV replication in LacDCs; rather, they further inhibited HIV replication in LacDCs without impacting cell viability (**Figure 4C**), indicating that extra cellular but not intracellular lactate may be involved in the induction of anti-HIV effect. To further characterize the function of lactate, fresh iDC was infected with HIV, and then cultured in the presence of 31.6 mM lactate and HIV replication was monitored. Lactate suppressed HIV replication in iDCs; therefore, lactate can inhibit HIV replication in differentiated iDC (**Figure 4D)**. An influx inhibitor, AZD3965, had no impact on HIV suppression in the presence of lactate but an influx and efflux inhibitor, Syrosingopine, partially restored the HIV inhibition (p<0.01) (**Figure 4D).** These results also suggested that extracellular, but not intracellular, lactate regulates HIV inhibition in DCs.

To determine whether lactate binds to cell-surface receptors/binding proteins and modifies gene expression, we compared gene expression profiles in iDC and lactate-treated iDC using bulk RNA-seq. To compare gene expression profiles, we cultured iDC with or without lactate for 1 day and 4 days from three independent donors’ cells. The volcano plot showed that lactate induced differentially expressed genes (DEGs), but no robust changes in anti-HIV genes expression [15,19] was observed (**Supplementary Figure 1)**; however, this result indicated that lactate stimulates cells through receptor(s). To define whether the well-described known lactate receptors (GPR81 and GRP132) express in iDC or LacDC, WB was conducted and expression was quantified (**Figure 4B and 4E)**. Both receptors were endogenously expressed in both cell types; GPR81 expression in LacDC increased by 4.4-fold compared with iDCs (n=4, p<0.05) **(Figure 4B**), whereas GPR132 was unchanged. In our previous studies, siRNA knockdown of the gene of interest was unsuccessful due to induction of cell death; therefore, in the current study, instead of knocking down GRP81 or GRP132, we compared DEGs using KEGG pathway [20] through the Database for Annotation, Visualization, and Integrated Discovery (DAVID) [21], because GRP81 and GRP132 signaling pathways have been demonstrated to be involved in cAMP/PKA, PI3K/AKT, and Yap/NF-κB, as well as ARRB2, PLC/IP3/Ca²⁺, ERK/STAT3/JAK2, and AKT/mTOR pathways in different cell types [8]. KEGG analysis showed that the PI3K/AKT signaling pathway ranked 35th on day 1 **(Supplemental Table 2)** but not on day 4 **(Supplemental Table 3)**. In the Day 4 DEGs, the TNF signaling pathway was ranked 24th, but other pathways were not listed. A volcano plot analysis was performed using those DEGs. The results demonstrated that lactate induced protein tyrosine kinase 2 (PTK2) on Day 1 (Supplemental Figure 2) and CCAAT enhancer-binding protein beta (CEBPB) and Fos proto-oncogene, AP-1 transcription factor subunit (FOS) on Day 4 **(Figure 4F),** respectively.

## Discussion

In our previous work, we investigated the mechanism of anti-HIV effect in 27DC; however, gene knock down analysis was unsuccessful in demonstrating a precise mechanism of anti-HIV [9]. This limitation is due to iDCs difficultly to transfect with siRNA or DNA and the transfection process itself can induce innate immune responses and cell death [22]. In the current study, we focused on metabolites due to a significant change in culture media and demonstrated that 27DC highly produces lactate, a pyruvate metabolite. This extracellular lactate can regulate HIV replication in DC; however, the lactate effect was not seen in macrophages, thus the effect is unique in DC. Further study will provide cell-type spectrum of the lactate effect.

Lactate exposure during differentiation reduced CD4 expression but not CCR5; consequently, virus binding, fusion, and RT reactions were suppressed accordingly. However, those steps showed no significant additional or synergistic suppression beyond the inhibition expected from reduced CD4, suggesting that the down regulation of CD4 plays the central role in resistance to HIV in LacDC. Although cell-surface CD4 expression was downregulated in LacDCs, total cellular CD4 protein levels were not significantly different from those in iDCs, suggesting that the reduction may involve altered receptor distribution in plasma membrane or intracellular comportment rather than degradation of the protein. Notably, CD4 transcript levels were significantly reduced in LacDCs despite unchanged total CD4 protein. This discrepancy may reflect the relatively longer half-life of CD4 protein stability in LacDC. Further studies are needed to determine whether altered receptor distribution, trafficking, internalization, or turnover contributes to reduced cell-surface CD4 expression in LacDCs.

Although lactate treatment elicits a comparatively limited number of DEGs in LacDC, it modified the expression profiles, suggesting that lactate may interact with cell-surface receptors or binding proteins and induce cell signaling. The major lactate receptors that have been identified are GRP81 and GRP132 [7]. As mentioned above, siRNA-mediated knockdown was unsuccessful in the DC study; to determine the receptor usage, we decided to predict receptors based on the gene expression profile using KEGG pathway analysis: GRP81 and GRP132 have been shown to induce the PI3K/AKT pathway and the PLC/Ca2 pathway, respectively [7,8]. KEGG pathway analysis indicated that some lactate-induced genes on the first day were associated with the PI3K/Akt pathway. The PI3K/Akt pathway has been demonstrated to play a pivotal role in the process of receptor internalization [23]. This observation suggests a potential mechanism by which lactate-mediated CD4 downregulation may occur through the GRP81/PI3/Akt pathway. Further studies are needed to identify the receptors used by lactate in iDCs. Additionally, the mechanism of downregulation in DCs and other cell types must be precisely defined.

## Conclusion

Lactate exposure during DC differentiation decreased cell-surface CD4 expression and suppressed HIV-1 infection. Lactate also suppressed HIV-1 replication in DCs that were already infected, indicating that it acts at a post-entry step in addition to limiting entry.

## Ethics Statement

Approval for this study, including all sample materials and protocols, was granted by the National Institute of Allergy and Infectious Diseases (NIAID) Institutional Review Board, and participants were provided the informed written consent prior to blood being drawn. All experimental procedures in these studies were approved by the National Cancer Institute at Frederick and Frederick National Laboratory for Cancer Research (the protocol code number: 16–19, approval date: 6 January 2017).

## Acknowledgement

The authors thank Veronique Nussenblatt for supporting this project. The content of this publication does not necessarily reflect the views or policies of the Department of Health and Human Services, nor does mentioning trade names, commercial products, or organizations imply endorsement by the U.S. Government. This project has been funded in whole or in part with federal funds from the National Cancer Institute, National Institutes of Health, under Contract No.75N91025F00012. This research was supported (in part) by the National Institute of Allergy and Infectious Diseases.

## Notes

### Competing Interest Statement

The authors have declared no competing interest.

## References

[1] L.A. O’Neill, R.J. Kishton, J. Rathmell, A guide to immunometabolism for immunologists, Nature reviews immunology 16 (2016) 553–565.

[2] B. Kelly, L.A. O’neill, Metabolic reprogramming in macrophages and dendritic cells in innate immunity, Cell research 25 (2015) 771–784.

[3] L. Wu, Z. Yan, Y. Jiang, Y. Chen, J. Du, L. Guo, J. Xu, Z. Luo, Y. Liu, Metabolic regulation of dendritic cell activation and immune function during inflammation, Frontiers in immunology 14 (2023) 1140749.

[4] E.J. Pearce, B. Everts, Dendritic cell metabolism, Nature Reviews Immunology 15 (2015) 18–29.

[5] S.Y. Lunt, M.G. Vander Heiden, Aerobic glycolysis: meeting the metabolic requirements of cell proliferation, Annual review of cell and developmental biology 27 (2011) 441–464.

[6] M.J. Rogatzki, B.S. Ferguson, M.L. Goodwin, L.B. Gladden, Lactate is always the end product of glycolysis, Frontiers in Neuroscience Volume 9–2015 (2015). 10.3389/fnins.2015.00022.

[7] M. Certo, A. Llibre, W. Lee, C. Mauro, Understanding lactate sensing and signalling, Trends in Endocrinology & Metabolism 33 (2022) 722–735.

[8] I. Manoharan, P.D. Prasad, M. Thangaraju, S. Manicassamy, Lactate-dependent regulation of immune responses by dendritic cells and macrophages, Frontiers in immunology 12 (2021) 691134.

[9] T. Imamichi, Q. Chen, B. Sowrirajan, J. Yang, S. Laverdure, M. Marquez, A.R. Mele, C. Watkins, J.W. Adelsberger, J. Higgins, Interleukin-27-induced HIV-resistant dendritic cells suppress reveres transcription following virus entry in an SPTBN1, autophagy, and YB-1 independent manner, Plos one 18 (2023) e0287829.

[10] Q. Chen, S. Swaminathan, D. Yang, L. Dai, H. Sui, J. Yang, R.L. Hornung, Y. Wang, D.W. Huang, X. Hu, Interleukin-27 is a potent inhibitor of cis HIV-1 replication in monocyte-derived dendritic cells via a type I interferon-independent pathway, PloS one 8 (2013) e59194.

[11] T.S. Theodore, G. Englund, A. Buckler-White, C.E. Buckler, M.A. Martin, K.W. Peden, Construction and characterization of a stable full-length macrophage-tropic HIV type 1 molecular clone that directs the production of high titers of progeny virions, AIDS research and human retroviruses 12 (1996) 191–194.

[12] M. Cavrois, C. De Noronha, W.C. Greene, A sensitive and specific enzyme-based assay detecting HIV-1 virion fusion in primary T lymphocytes, Nat Biotechnol 20 (2002) 1151–1154. 10.1038/nbt745.

[13] M. Cavrois, J. Neidleman, W. Yonemoto, D. Fenard, W.C. Greene, HIV-1 virion fusion assay: uncoating not required and no effect of Nef on fusion, Virology 328 (2004) 36–44.

[14] T. Imamichi, S.C. Berg, H. Imamichi, J.C. Lopez, J.A. Metcalf, J. Falloon, H.C. Lane, Relative replication fitness of a high-level 3’-azido-3’-deoxythymidine-resistant variant of human immunodeficiency virus type 1 possessing an amino acid deletion at codon 67 and a novel substitution (Thr-->Gly) at codon 69, J Virol 74 (2000) 10958–10964. 10.1128/jvi.74.23.10958-10964.2000.

[15] T. Imamichi, J. Yang, Q. Chen, S. Goswami, M. Marquez, U. Kariyawasam, H.N. Sharma, R. Wiscovitch-Russo, X. Li, A. Aioi, Interleukin-27-polarized HIV-resistant M2 macrophages are a novel subtype of macrophages that express distinct antiviral gene profiles in individual cells: implication for the antiviral effect via different mechanisms in the individual cell-dependent manner, Frontiers in immunology 16 (2025) 1550699.

[16] L. Dai, K.B. Lidie, Q. Chen, J.W. Adelsberger, X. Zheng, D. Huang, J. Yang, R.A. Lempicki, T. Rehman, R.L. Dewar, Y. Wang, R.L. Hornung, K.A. Canizales, S.J. Lockett, H.C. Lane, T. Imamichi, IL-27 inhibits HIV-1 infection in human macrophages by down-regulating host factor SPTBN1 during monocyte to macrophage differentiation, J Exp Med 210 (2013) 517–534. 10.1084/jem.20120572.

[17] T. Nishi, M. Forgac, The vacuolar (H+)-ATPases—nature’s most versatile proton pumps, Nature reviews Molecular cell biology 3 (2002) 94–103.

[18] A.P. Halestrap, The monocarboxylate transporter family—structure and functional characterization, IUBMB life 64 (2012) 1–9.

[19] T. Imamichi, J. Yang, Q. Chen, U. Kariyawasam, M. Marquez, J. Higgins, J. Metz, H.N. Sharma, M.W. Baseler, H. Sui, Integrated Functional and scRNA-Seq Analyses Reveal Convergence of M-CSF- and GM-CSF-Derived Macrophages Following IL-27 Polarization, Cells 15 (2026). 10.3390/cells15060528.

[20] H. Ogata, S. Goto, K. Sato, W. Fujibuchi, H. Bono, M. Kanehisa, KEGG: Kyoto encyclopedia of genes and genomes, Nucleic acids research 27 (1999) 29–34.

[21] B.T. Sherman, M. Hao, J. Qiu, X. Jiao, M.W. Baseler, H.C. Lane, T. Imamichi, W. Chang, DAVID: a web server for functional enrichment analysis and functional annotation of gene lists (2021 update), Nucleic Acids Res 50 (2022) W216–w221. 10.1093/nar/gkac194.

[22] O. Gresch, L. Altrogge, Transfection of difficult-to-transfect primary mammalian cells, Protein Expression in Mammalian Cells: Methods and Protocols, Springer 2011, pp. 65–74.

[23] P.-h. Chen, H. Yao, L.J.-s. Huang, Cytokine receptor endocytosis: new kinase activity-dependent and-independent roles of PI3K, Frontiers in Endocrinology 8 (2017) 78.

