## Supplemental method for "Extracellular Lactate Inhibits HIV Infection in Dendritic Cells by Downregulating CD4 Expression"

**Supplemental methods**

**Bulk RNAseq data analysis**

Raw sequencing data in Fastq format were processed using Partek Flow software (https://www.illumina.com/products/by-type/informatics-products/partek-flow.html). Reads were mapped to the human reference genome (hg38) **(https://www.ncbi.nlm.nih.gov/datasets/genome/GCF_000001405.26/** ) using the HISAT2 alignment algorithm. Transcript quantification and gene-level summarization were performed using the Ensembl database (**https://www.ensembl.org/Homo_sapiens/Info/Index**). Following alignment, gene expression levels were normalized using the trimmed mean of M-values (TMM) method [11]. To identify differentially expressed genes (DEGs), an ANOVA was employed, incorporating Donor as a random effect and an interaction term for Cell Type and Treatment. Significant DEGs were defined as those exhibiting fold change ≥2.and  *p*-values < 0.05. To visualize the overall distribution of differentially expressed genes, a two-dimensional volcano plot was generated within Partek Flow (Partek Inc., St. Louis, MO, USA). Following ANOVA analysis, the volcano plot was invoked directly from the statistical results spreadsheet using the visualization menu. Genes were mapped with log2 fold change plotted on the x-axis and -log10p-value plotted on the y-axis.
