## supplemental figure1 for "Extracellular Lactate Inhibits HIV Infection in Dendritic Cells by Downregulating CD4 Expression"

### Slide 1
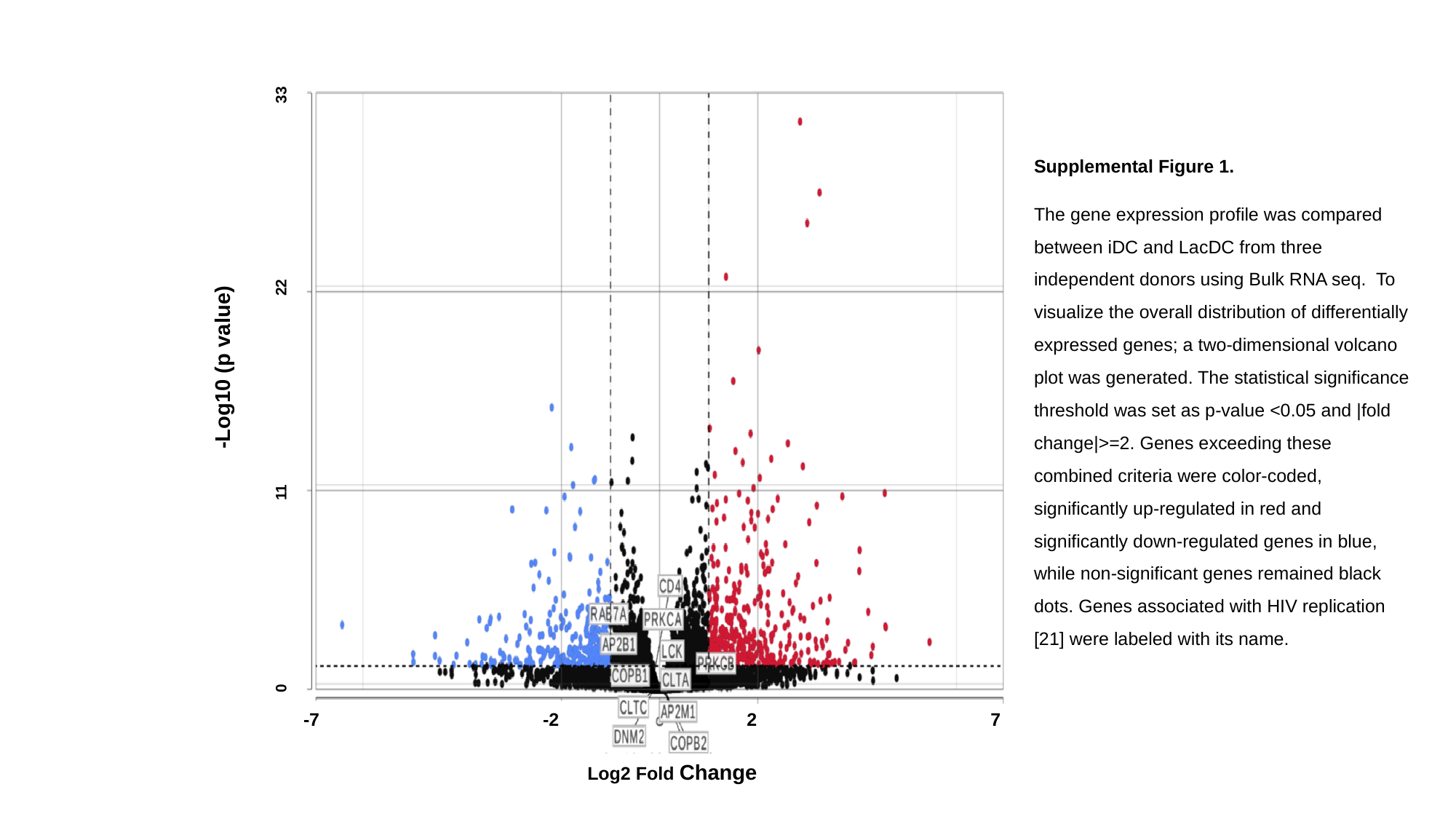

-Log10 (p value)
0 11 22 33
-7 -2 2 7
Log2 Fold Change
Supplemental Figure 1.
The gene expression profile was compared between iDC and LacDC from three independent donors using Bulk RNA seq. To visualize the overall distribution of differentially expressed genes; a two-dimensional volcano plot was generated. The statistical significance threshold was set as p-value <0.05 and |fold change|>=2. Genes exceeding these combined criteria were color-coded, significantly up-regulated in red and significantly down-regulated genes in blue, while non-significant genes remained black dots. Genes associated with HIV replication [21] were labeled with its name.

### Slide 2
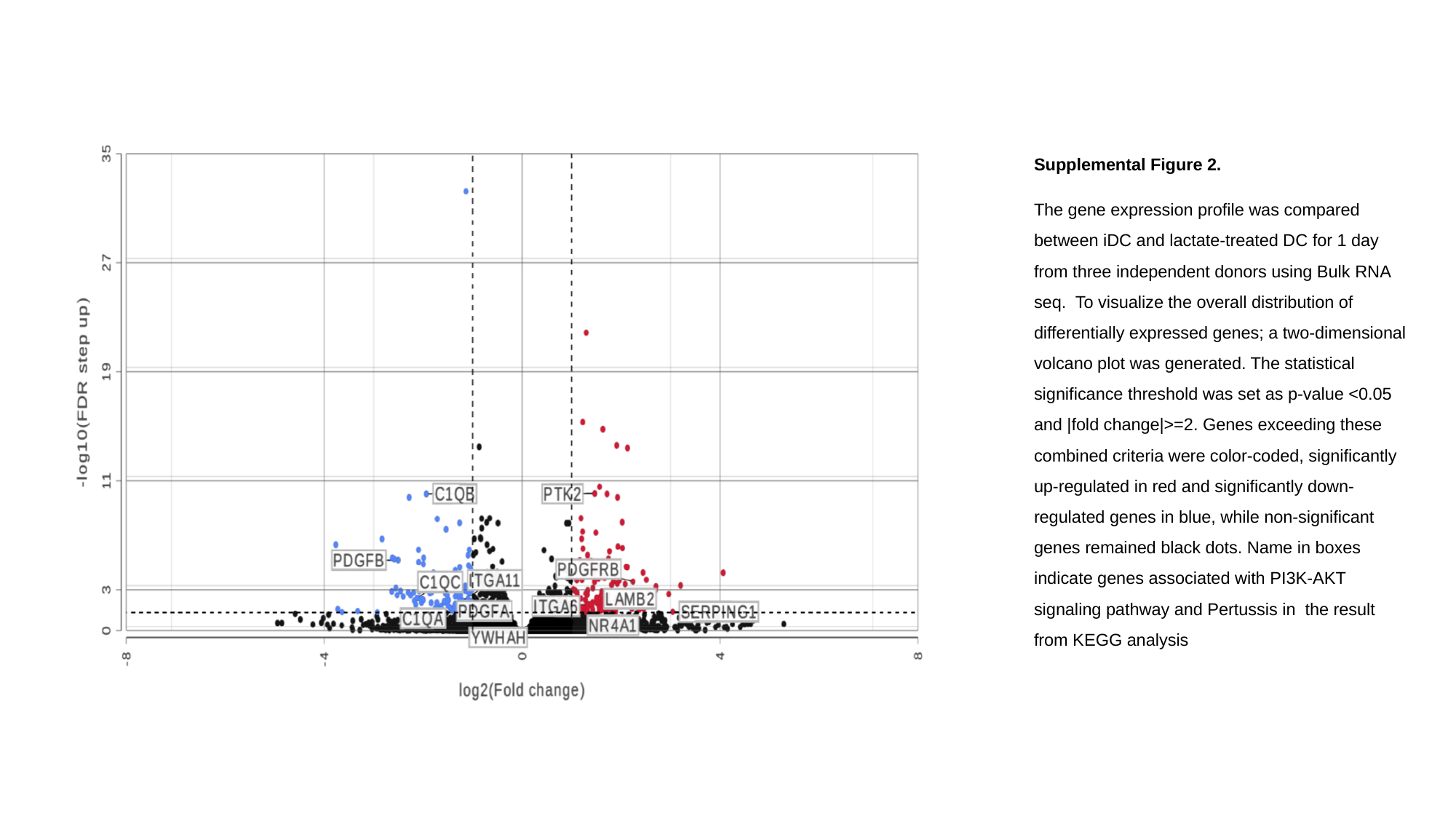

Supplemental Figure 2.
The gene expression profile was compared between iDC and lactate-treated DC for 1 day from three independent donors using Bulk RNA seq. To visualize the overall distribution of differentially expressed genes; a two-dimensional volcano plot was generated. The statistical significance threshold was set as p-value <0.05 and |fold change|>=2. Genes exceeding these combined criteria were color-coded, significantly up-regulated in red and significantly down-regulated genes in blue, while non-significant genes remained black dots. Name in boxes indicate genes associated with PI3K-AKT signaling pathway and Pertussis in the result from KEGG analysis
